# The AMPKα2/PHF2 axis is critical for turning over lipid droplets during muscle stem cell fate

**DOI:** 10.1101/2025.01.18.630727

**Authors:** Delia Cicciarello, Sandrine Mouradian, Mélodie Pitchecanin, William Jarassier, Anita Kneppers, Rémi Mounier, Laurent Schaeffer, Isabella Scionti

## Abstract

Lipid metabolism is a key process required for muscle stem cell (MuSCs) function during regenerative myogenesis. However, the molecular pathway responsible of such regulation is still unknown. Studies of lysine demethylase PHF2 have reported its critical role in lipid metabolism in pathological processes, although there are no data regarding its role during MuSC fate transition. Here we show that PHF2 controls lipid droplet homeostasis in MuSCs during regenerative myogenesis, by promoting the contact between lipid droplets and mitochondria. Consistently, in absence of PHF2, myocytes accumulate lipid droplets, leading to mitochondrial dysfunction and impaired regeneration. Interestingly, such phenotype is rescued by AMPKα2-PHF2 phospho-mimetic mutant expression. Our findings provide evidence that PHF2 is part of AMPKα2 signaling and underscore the critical role of AMPKα2/PHF2 axis in regulating lipid droplets homeostasis during MuSC fate.

## Introduction

Cellular metabolism has emerged as a key regulator of adult stem cell function supporting tissue maintenance and regeneration. Indeed, in the last decade it has been shown that stem cell fate is characterized by a specific shift in energy metabolic profiles that are common to different adult stem cells^1^. While the importance of glycolysis and fatty acid oxidation (FAO) in the maintenance respectively of stem cell proliferation and quiescence is recognized^2,3^, there is limited understanding of how processes like lipogenesis, lipolysis, and lipophagy regulate stem cell fate. Cells have the capacity to store lipids in the form of lipid droplets (LD)^4^. For a long time, LDs have been considered as inert cytosolic lipid storage organelles and have only recently been recognized as dynamic and crucial actors of lipid uptake, metabolism, trafficking, and signaling in the cell^5^. Indeed, LDs are dynamically synthesized and catabolized in response to cellular needs and environmental signals. Besides LD role in the development of diseases, such as non-alcoholic fatty liver disease (NAFLD), obesity and Tauopathy^6–8^, the mechanisms underlying the turnover of LDs and their biological importance in stem cell fate have remained largely unexplored.

Adult skeletal muscle has the unique ability to regenerate after an injury or following a trauma, relying on the presence of muscle stem cells (MuSCs). While MuSCs are quiescent in unperturbed muscle, they activate upon muscle injury, proliferate and then commit to the myogenic fate by differentiating and fusing into muscle fibers (myofibers). A subset of activated MuSCs returns to quiescence (self-renewal) to replenish the pool of MuSCs^9^. MuSCs, like all other cells, require energy (ATP) to carry out the reactions necessary for their functions. However, the metabolic demands differ among quiescent MuSCs, activated/proliferating myoblasts and differentiating myocytes. Accordingly, it was reported that MuSC fate is supported by a profound alterations in metabolic pathways, called metabolic reprogramming^10–12^. Recently, it has been described that LD abundance and turnover influence MuSC fate^13^. Quiescent and self-renewed MuSCs are characterized by low LD content, since their metabolism mostly relies on FAO^2,13^. Upon activation LDs accumulate and reach their highest concentration in committed myocytes^13^. Later, LDs need to be catabolized to allow the fusion of committed myocytes into multinucleated myotubes^13^. As many pathways involved in lipid metabolism are not localized to LDs, close connection with other cell organelles is required^14^. While the biogenesis of LDs takes place within the Endoplasmic reticulum (ER) membrane^15^; Following nutrient stress, the two cellular energy sensors, cAMP-dependent protein kinase A (PKA) and the AMP-activated protein kinase (AMPK)^16^, trigger LD catabolism^17^ in order to release fatty acids (FAs) as an energy source. Since energy production from fatty acids occurs in the mitochondria, the physical and functional interplay between lipid droplets and mitochondria is essential for the initiation of LD catabolism^18^.

PHD-finger protein 2 (PHF2/KDM7C) demethylase belongs to the lysine demethylase 7 (KDM7) family of histone H3 lysine 9 (H3K9), H3K27 and H4K20 demethylases. PHF2 protein contains a plant homeodomain (PHD), involved in protein/protein or protein/nucleic acid interactions, and a Jumonji (JmjC) domain, which, in contrast to the other lysine demethylases, does not have enzymatic activity by itself^19^. Indeed PHF2 demethylase activity requires activation through post-translational phosphorylation by the two energetic sensors PKA and AMPK^20,21^, in order to demethylate its protein targets. This unique mode of activation suggests a specific regulatory mechanism for PHF2 upon energy stress, supporting the hypothesis that PHF2 enzymatic activity is necessary for MuSC metabolic adaptation during skeletal muscle regeneration.

Here we demonstrate that PHF2, as a component of the AMPKα2 metabolic signaling pathway, plays a crucial role in maintaining energy homeostasis in MuSCs during regenerative myogenesis, particularly by modulating LD-mitochondria interaction.

## Results

### PHF2 depletion impairs skeletal regenerative myogenesis

To assess the physiological role of PHF2 *in vivo* during regenerative myogenesis, PHF2^fl/fl22^ mice were crossed with Pax7^CreERT2^ mice^23^ (hereafter named **PHF2 SCiKO**). Nine-week-old mice were treated with tamoxifen (TAM) and all MuSCs (PAX7^pos^) are knocked-out for PHF2 (**Figure S1A**). One week after the last TAM injection, *Tibialis anterior* (TA) muscles were analyzed before and at several time points after cardiotoxin (CTX) injury (**Figure S1B**). In PHF2 SCiKO muscle, histological features (**Figure S1C**) as well as the number and size of fibers per mm^2^ were unaffected compared to CTRL SC mice before injury (**Figure S1D, E**) and 7-days post injury (dpi) (**Figure S1F, G**). However, 11 and 28 dpi, PHF2 SCiKO TA muscles exhibited smaller fibers compared to CTRL SC muscles as evidenced by the cross-sectional area analysis (**Figure 1A-D**). Moreover, despite the equal number of quiescent PAX7^pos^ cells per mm^2^ in uninjured condition between the two genotypes (**Figure S1H**) and the percentage of proliferating myoblasts (PAX7^pos^KI67^pos^) at 7 dpi (**Figure S1I**), we observed a significant decreased number of MuSCs that have returned to quiescence at 28 dpi (PAX7^pos^/KI67^neg^) (**Figure 1E**). We next quantified the number of MYOG^pos^ cells at 11 dpi and we observed a significantly increased number of MYOG^pos^ cells, which likely failed to fuse, in PHF2 SCiKO muscles (+52%) (**Figure 1F**). These results suggest that PHF2 is necessary for MuSCs function *in vivo* upon injury, since PHF2 depletion results into impairment of MuSC regenerative and self-renewal potential.

**Figure 1.**
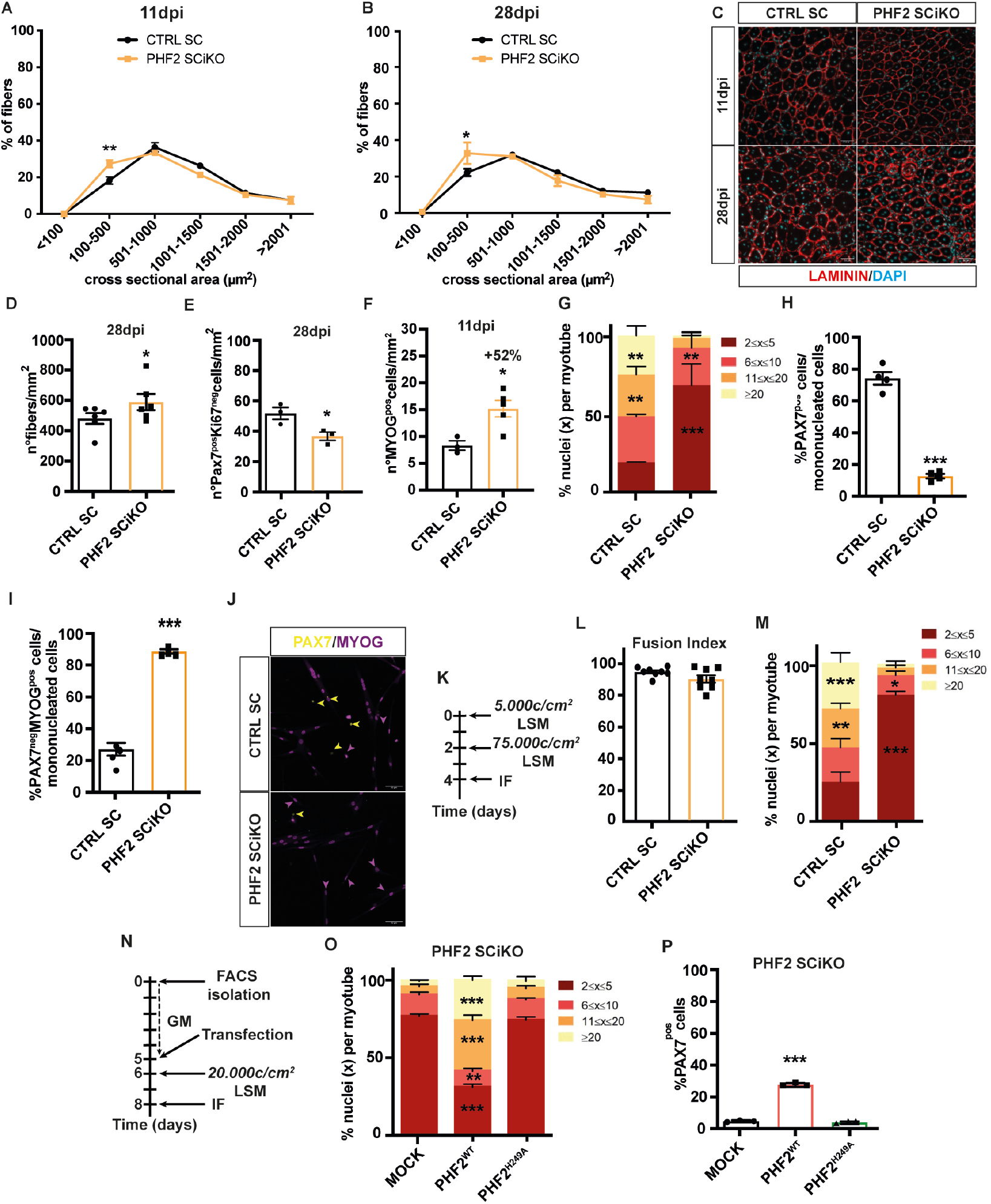
PHF2 ablation impairs regenerative myogenesis. CSA distribution of muscle fibers on cryosections of regenerated TA muscles at 11dpi (A) and 28dpi (B). (C) Representative images of immunostainings. (D) Number of myofibers per mm2 at 28dpi. (E) Quantification of the number of PAX7^pos^Ki67^neg^ cells per mm^2^ at 28 dpi. (F) Quantification of the number of MYOG^pos^ cells per mm^2^ on CTRL SC and PHF2 SCiKO TA cryosections at 11dpi. (G) Percentage of nuclei per myotube, (H) Percentage of PAX7^pos^MYOG^neg^ (yellow arrowhead) and (I) PAX7^neg^MYOG^pos^ cells (purple arrowhead) in cultures after 48h in LSM. (J) Representative images of immunostainings. (K) Experimental setup. Fusion index (L) and percentage of nuclei per myotube (M) in cultures after 48h in LSM. (N) Experimental setup. Percentage of nuclei per myotube (O) and of PAX7^pos^ cells (P) in PHF2 SCiKO cells transfected with either MOCK, PHF2^wt^ or PHF2^H249A^. Scale bars, 50 μm. n = 3-6 mice/genotype. n = 3-8 primary cultures/genotype. Values are mean or percentage mean ± SEM. *P < 0.05, **P < 0.01, ***P < 0.001 (Student *t*-test [D-E-F-H-I-L panels], Sidak’s test after two-way ANOVA [A-B panels], Tukey’s test after one way [G-M-P panels] or two-way ANOVA [O panel]).

We next dissected the role of PHF2 in adult MuSC fate *in vitro*. MuSCs were fluorescence activated cell sorting (FACS)-isolated using a specific cocktail of surface antibodies, amplified, and then induced to fully differentiate into low serum medium (LSM) (**Figure S1J**). Two days after, cells were labeled for PAX7 (mononucleated self-renewing MuSCs), MYOG (mononucleated myocytes) and MYHC (multinucleated myotubes) expression. While CTRL SC and PHF2 SCiKO cells had identical growth rates (**Figure S1K**) and similar fusion index (**Figure S1L**), 73% of PHF2 SCiKO myotubes contained only 2 to 5 nuclei compared to 21% of control myotubes (**Figure 1G**), supporting that PHF2 plays a role in the MuSC differentiation process. Moreover, while the percentage of mononucleated cells (26% in CTRL SC and 26% in PHF2 SCiKO) was similar (**Figure S1M)**, PHF2 SCiKO cultures showed a significantly decreased rate (12%) of PAX7^pos^ self-renewed MuSCs compared to CTRL SC cultures (74%) (**Figure 1H**). Interestingly the majority of PHF2 SCiKO mononucleated cells were PAX7^neg^MYOG^pos^ (88%) compared to the CTRL SC cells (26%), suggesting a defect in the fusion process (**Figure 1I, J**). To test this hypothesis, freshly isolated MuSCs were seeded in LSM at low density and let to differentiate into myocytes (**Figure 1K and Figure S1N**)^24^. Differentiated myocytes were then plated at high density in LSM to evaluate their fusion capacity (**Figure 1K**)^24^. While the fusion index was identical in PHF2 SCiKO and CTRL cultures (**Figure 1L)**, only 20% of PHF2 SCiKO myotubes contained more than 5 nuclei compared to 75% of CTRL SC myotubes, including 30% of myotubes containing more than 20 nuclei (**Figure 1M**). To ensure that differentiation defect was due to PHF2 enzymatic activity, rescue experiments were performed by expressing either a human *wild-type* PHF2 (PHF2^wt^) or a catalytically inactive PHF2 mutant (PHF2^H249A^)^20^ (**Figure 1N**). Over-expression of PHF2^wt^ efficiently restored the myotube formation (**Figure 1O**) and self-renewal capacity (**Figure 1P**) in PHF2 SCiKO cells compared to Mock cells. Conversely, the PHF2^H249A^ mutant failed to rescue the phenotype (**Figure 1O, P**). These results demonstrate the requirement of PHF2 and of its de-methylase activity for MuSC fusion potential.

### PHF2 inactivation causes accumulation of lipid droplets in committed myocytes

Perturbation of lipid droplet biogenesis or catabolism severely impairs MuSCs differentiation potential^13^. In the context of pathological conditions, the loss of PHF2 leads to LD accumulation^25,26^. To determine whether PHF2 could influence LD homeostasis during regenerative myogenesis, we performed a time course of stage-specific marker expression during proliferation and differentiation of MuSCs freshly isolated from CTRL and PHF2 SCiKO muscles. Cells were labelled with PAX7, MYOD (commitment into myogenic lineage), MYOG and BODIPY (neutral lipid probe) in proliferation (growth medium, GM) and at different time-points (12h, 24h, 36h, 48h) during differentiation in LSM. Similar percentages of proliferating myoblasts (PAX7^pos^MYOD^pos^), committed myoblasts (PAX7^neg^MYOD^pos^) as well as differentiated myocytes (PAX7^neg^MYOG^pos^), which accumulate lipid droplets (BODIPY^pos^) were observed in GM and after 12h and 24h in LSM (**Figure S2A-F**). Conversely, at 36h and 48h, the percentage of differentiated myocytes (PAX7^neg^MYOG^pos^, 80,33% at 36h and 87% at 48h), accumulating lipid droplets (PAX7^neg^MYOG^pos^BODIPY^pos^, 66% at 36h and 86% at 48h) was significantly higher in PHF2 SCiKO cultures compared to CTRL SC ones (**Figure S2D-F and Figure 2 A, B**). These results demonstrate a link between PHF2, lipid droplet metabolism and myocyte fate. To further confirm this observation *in vivo*, cryosections of 11dpi injured TA muscles were stained for MYOG and Perilipin, a LD membrane-associated proteins. A significant accumulation of MYOG^pos^Perilipin^pos^ cells (16%) was observed in PHF2 SCiKO regenerating muscles compared to CTRL SC (4%, **Figure 2C-D**). Previous findings^25^, in the context of cancer, have shown that PHF2 loss leads to lipid droplet accumulation by activating lipogenesis through regulation of SREBP1 protein stability. However, in PHF2 SCiKO MYOG^pos^ myocytes we observed an increase neither in SREBP1 protein nor in DGAT2 protein related to lipogenesis (**Figure S2G, H**). These results support the idea that in muscle cells PHF2 inactivation causes LD accumulation via other mechanisms.

**Figure 2.**
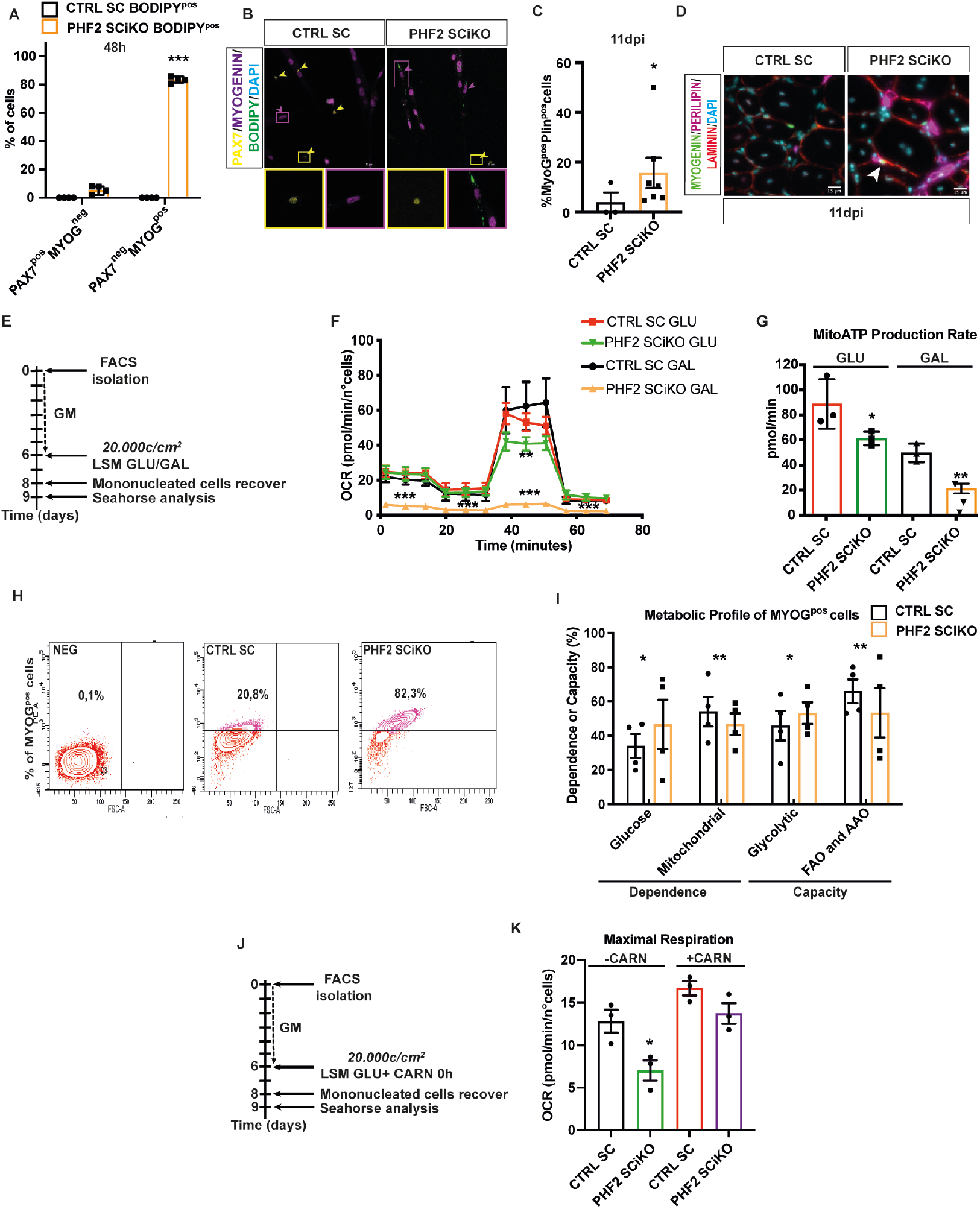
PHF2 KO myocytes accumulate LDs displaying metabolic defects. (A) Percentage of PAX7^pos^MYOG^neg^ BODIPY^pos^ (yellow arrowhead) and PAX7^neg^MYOG^pos^BODIPY^pos^ cells (purple arrowhead) after 48h in LSM. (B) Representative images. (c)Percentage of MYOG^pos^Perilipin^pos^ cells on cryosections of regenerated TA muscles at 11 dpi. (D) Representative images (MYOG^pos^Perilipin^pos^ cell is indicated by white arrowhead). (E) Experimental setup. (F) Oxygen consumption rate (OCR) and (G) mitochondrial ATP rate of mononucleated cells. Percentage of MYOG^pos^ cells (H) and Metabolic profile (I) of MYOG^pos^ cells after 48h in LSM. (J) Experimental setup. (K) Maximal respiration measure after L-carnitine supplementation. Scale bars, 15 and 50 μm. n = 3 mice/genotype. n = 3-4 primary MuSC cultures/genotype. Values are mean or percentage mean ± SEM. *P < 0.05, **P < 0.01, ***P < 0.001 (Student *t*-test [A-C-G-I panels] and Tukey’s test after one way-ANOVA [F-K panels]).

### PHF2 KO myocytes displayed a different metabolic profile than control cells

Defects in lipid metabolism are associated with metabolic alterations^*27*^. Thus, we measured the mitochondrial respiratory capacity of mononucleated CTRL and PHF2 KO cells after 48h in LSM, using a Seahorse analyzer (**Figure 2E**). In absence of PHF2 we observed a significant decrease in the mitochondrial maximal respiratory rate (**Figure 2F**) together with reduced ATP production (**Figure 2G**). In order to test whether the observed dysfunction was linked to perturbation of FAO, we differentiated CTRL and PHF2 KO cells in LSM supplemented with galactose (GAL) instead of glucose (GLU) (**Figure 2E**). Oxidation of galactose to pyruvate through glycolysis yields no net production of ATP thus forcing cells to rely more on FAs and amino acids in order to produce ATP^*28*^. Strikingly, PHF2 KO cells, differentiated in GAL, displayed a severe impairment in their mitochondrial capacity to produce ATP by using FAs or amino acids, as shown by a marked reduction of both basal and maximal respiration (**Figure 2F, G**), suggesting that PHF2 KO cells rely more on glucose as source of energy and are incapable of undergoing FAO. To further confirm this hypothesis, we used a novel flow cytometry-based method (SCENITH)^*29*^ that allows to determine energy metabolism at single cell level. Consistently, PHF2 KO MYOG^pos^ cells (**Figure 2H)** showed an increased reliance on glucose compared to CTRL cells (**Figure 2I**), supporting the conclusion that these cells have an impairment in the metabolic reprogramming toward FAO. Furthermore, to decipher whether the mitochondrial FAO dysfunction of PHF2 KO myocytes was due to absence of FA substrates or to the inability to metabolize them, mitochondrial oxygen consumption rate (OCR) was measured after L-carnitine supplementation (**Figure 2J)**. Strikingly, OCR was entirely rescued in PHF2 SCiKO cells treated with L-Carnitine compared to non-treated PHF2 SCiKO cells (**Figure 2K)**. These findings provide evidence that the mitochondrial dysfunction observed in PHF2 KO cells is attributable to the lack of fatty acid availability rather than to an intrinsic mitochondrial inability to metabolize them, supporting the hypothesis that PHF2 is involved in lipid metabolism or in the FA trafficking into the mitochondria.

### AMPKα2-mediated phosphorylation of PHF2 is required for lipid droplet turnover in myocytes

To decipher to which metabolic pathway PHF2 belongs, rescue experiments were performed by over-expressing either a human *wild-type* PHF2 (PHF2^wt^), or catalytically inactive PHF2 (PHF2^H249A^) or phospho-mimetic mutants of PHF2 by AMPKα2 (PHF2^S655E^) or PKA (PHF2^S1056E^) or PHF2 mutants non-phosphorilatable by AMPKα2 (PHF2^S655A^) or PKA (PHF2^S1056A^). Expression of PHF2^wt^ and AMPKα2-phospho-mimetic PHF2^S655E^ completely rescued the LD accumulation phenotype in PHF2 KO myocytes (**Figure 3A-B**). Conversely, the PHF2^H249A^, AMPKα2-PHF2^S655A^ and PKA-PHF2^S1056E/S1056A^ did not prevent LD accumulation (**Figure 3A-B and Figure S3A-B**). Since the rate of AMPK phosphorylation was similar between CTRL and PHF2 KO myocytes (**Figure S3C-D**), these data demonstrate that PHF2 phosphorylation on S655 is necessary for lipid droplet turnover. To further confirm this conclusion, we took advantage of a mouse model lacking functional AMPKα2 (AMPKα2 ^-/-^)^30^. MuSCs were FACS-isolated from wild-type (WT) and AMPKα2 ^-/-^ mice, seeded and fully differentiated *in vitro* (**Figure S1J**). As previously reported, ablation of AMPKα2 impaired myocytes fusion (data not shown)^30^. In addition, we observed a significantly increased percentage of mononucleated MYOG^pos^ cells that accumulate lipid droplets in AMPKα2 ^-/-^ cultures compared to *Wild-Type* (WT, **Figure 3C-E**). Strikingly, by overexpressing AMPKα2-PHF2 phospho-mimetic (PHF2^S655E^) mutant, lipid droplet accumulation in MYOG^pos^ cells was completely rescued in AMPKα2 ^-/-^ cultures (**Figure 3F, Figure S3E**). Collectively these results demonstrate that the AMPKα2/PHF2 axis is required for LD turnover.

**Figure 3.**
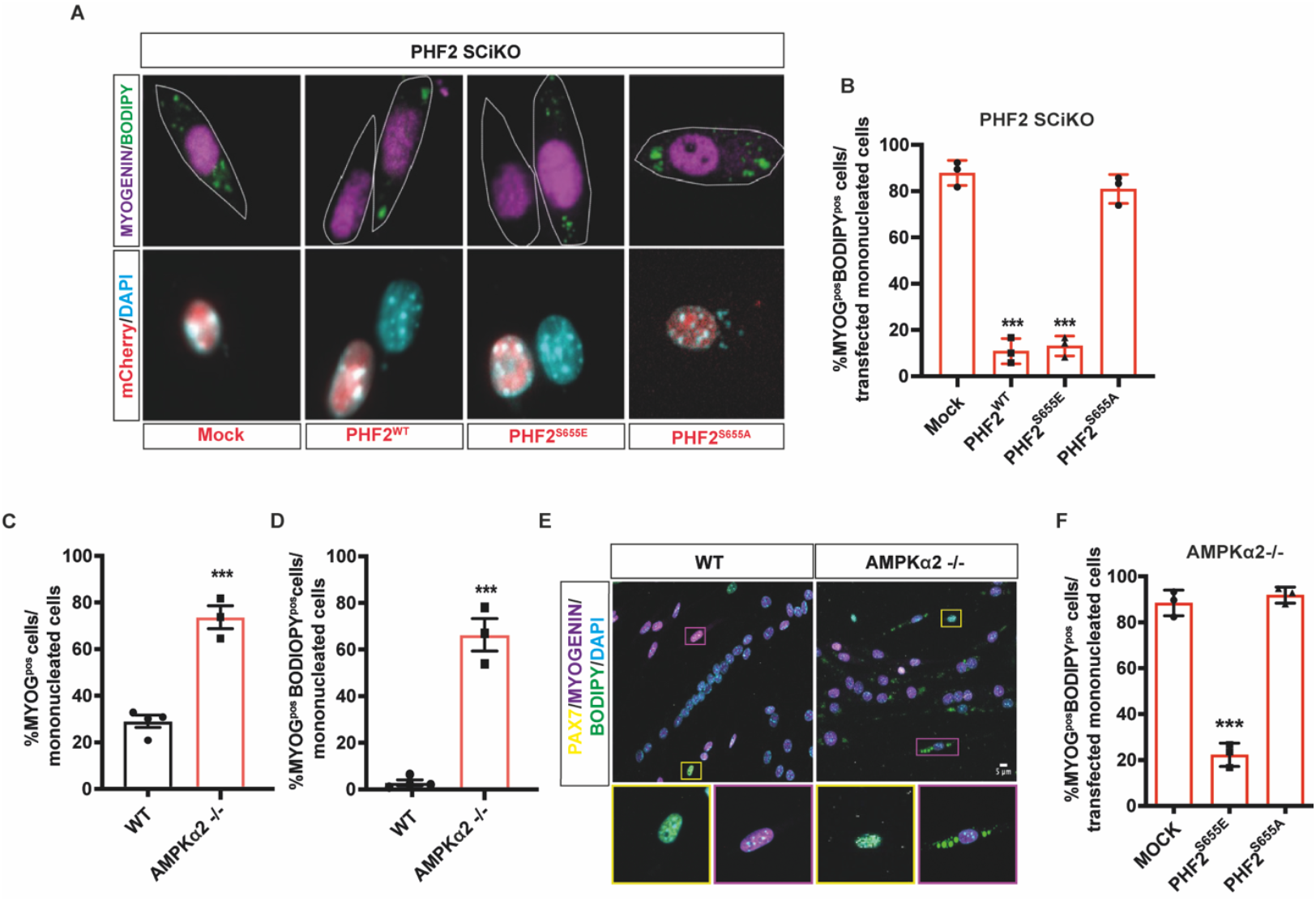
AMPKα2 mediated-activation of PHF2 regulates LD turnover. (A-B) Percentage of MYOG^pos^BODIPY^pos^ cells transfected with either MOCK, PHF2^wt^ or PHF2^S655E^ and PHF2^S655A^. Representative images. (C) Percentage of MYOG^pos^ and (D) percentage of MYOG^pos^BODIPY^pos^ cells after 48h in LSM. (E) Representative images. (F) Percentage of MYOG^pos^BODIPY^pos^ cells transfected with either MOCK, PHF2^S655E^ or PHF2^S655A^. Scale bars, 5 μm. n = 3 primary cultures/genotype. Values are mean or percentage mean ± SEM. ***P < 0.001 (Student *t*-test [C-D panels] and Tukey’s test after one way-ANOVA [B-F panels]).

### PHF2 mediates LD metabolism by regulating the interaction between LDs and mitochondria

To explore the molecular mechanism underlying LD accumulation, we used a *Pax7* ^*CreERT2*^; *R26R*^*EYFP/EYFP*^; *PHF2*^*flox/flox*^ model, that expresses EYFP in PHF2 KO MuSCs and their descendants. We performed a single-cell RNA sequencing (scRNA-seq) on FACS-isolated EYFP^pos^ cells at 11dpi from CTRL SC and PHF2 SCiKO mice. Unsupervised clustering of the quality-controlled cells returned 5 cell clusters, transcriptomes of which were related to MuSCs (*Pax7*^*pos*^ and *Notch3*^*pos*^) (**Figure 4A, Figure S4A**), proliferating myoblasts (*Pax7*^*pos*^, *Myod1*^*pos*^ and *Mki67*^*pos*^) (**Figure 4A, Figure S4B**), myocytes (*Myog*^*pos*^), and myocytes undergoing fusion, as evidenced by the co-expression of early myosin isoforms (*Myog*^*pos*^, *Myh3*^*pos*^, *Myh8*^*pos*^) (**Figure 4A, B**), Myofibers, and ‘Others’ (**Figure 4A)**. We then analyzed the lipid droplet metabolic pathways, and consistently with the observed phenotype, LD membrane related genes *Plin2* and *Plin3* were upregulated in the PHF2 SCiKO myocyte cluster compared to CTRL SC (**Figure 4C**). However, the expression of genes annotated with the gene ontology (GO) terms for lipolysis (GO:0046461), lypophagy (GO:0061724) or LD biogenesis (GO:0019432) processes were comparable between the two conditions (**Figure S4C**).

**Figure 4.**
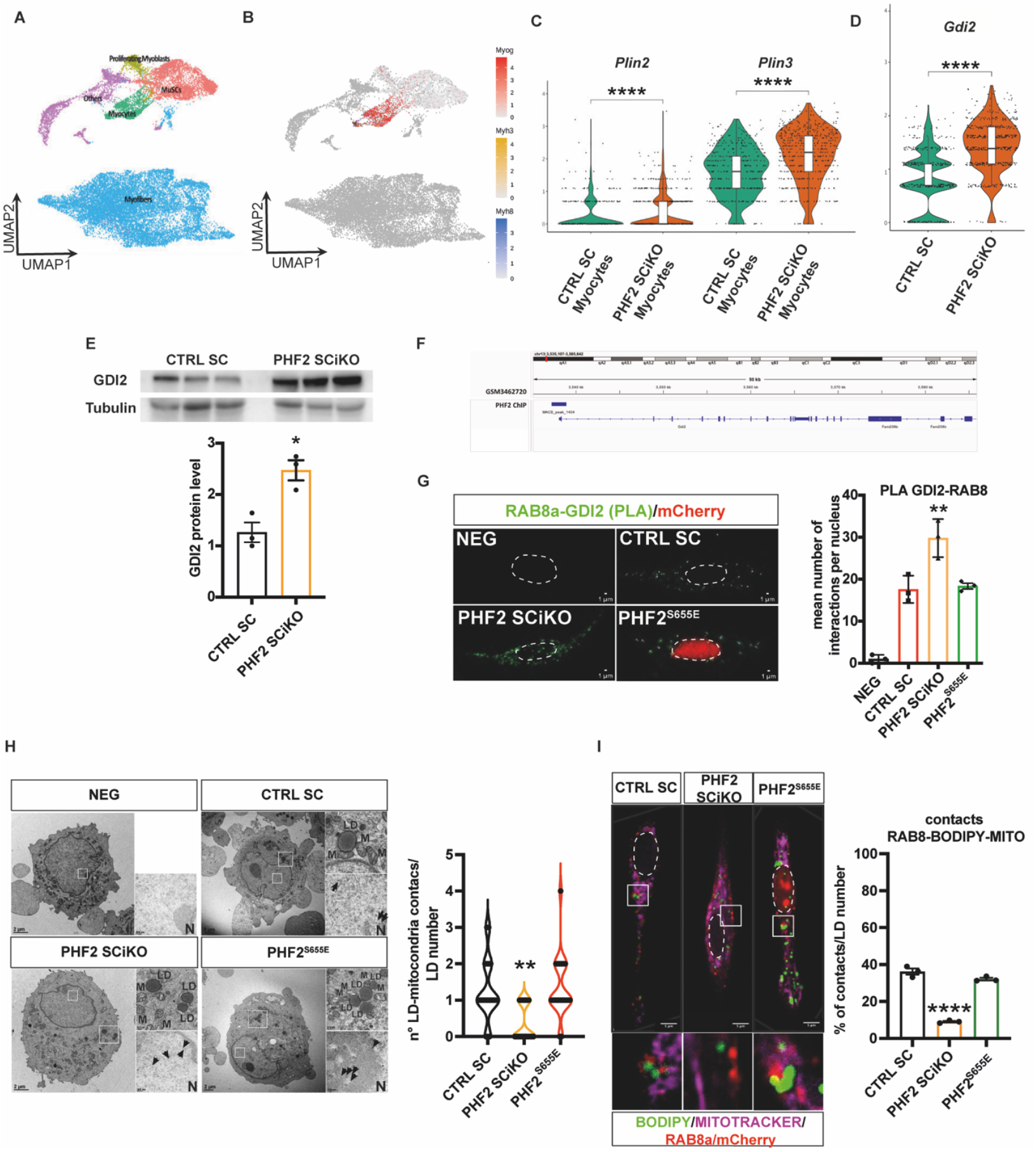
PHF2 regulates LD-mitochondria interactions. (A) Clustering of EYFP^pos^ -FACS-isolated cells from CTRL SC and PHF2 SCiKO at 11dpi (n=3 mice pooled in 1 sample/genotype). (B) *Myog*^*pos*^ myocytes cluster. *Plin2, Plin3* (C) and *Gdi2* (D) mRNA expression levels. (E) GDI2 protein level in CTRL SC and PHF2 SCiKO mononucleated cells 24h post LSM. (F) Integrative Genomics Viewer (IGV) snapshot depicting PHF2 enrichment (MACS peak call in dark blue) on the *Gdi2* promoter region (GSM3462720). PLA analysis (G), electron microscopy analysis of LD-mitochondria contacts (H) and confocal analysis of LD-RAB8a-mitochondria contacts (I) in CTRL SC and PHF2 SCiKO mononucleated cells 24h post LSM. Scale bars, 1, 2 and 5 μm. n = 3 primary cultures/genotype. For (H panel), n=20 cells/genotype/conditions. Values are mean or percentage mean ± SEM. **P < 0.01, ****P < 0.0001 (Student *t*-test [E panel] and Tukey’s multiple comparison test after one way-ANOVA [G-H-I panels]).

Lipid droplet metabolism involves dynamic interactions between LDs and mitochondria^31^, which are under the control of RAB8a protein^32^. RAB8a protein, like other small GTPases, cycles between the active GTP-bound and inactive GDP-bound forms, where the GTP-bound form confers biological activity. Such cycle is tightly regulated by guanine nucleotide exchange factors (GEFs), which catalyze the exchange from GDP to GTP; GTPase-activating proteins (GAPs), which switch off the signaling by inducing GTP hydrolysis; and GDP dissociation inhibitor proteins (GDIs), which binds to the GDP form and prevents the GDP to GTP exchange, keeping RAB8a soluble in the cytosol in its inactive form^33^. Violin plot reveals that the expression of *Rab8a*, GEF genes (*Rab3ip, Rab3il1, Rabif and C9orf72*) or GAP genes (*Tbc1d1* and *Tbc1d4*) was comparable between the two genotypes or barely detectable (**Figure S4D-F**). However, the expression of the ubiquitous *Gdi2* gene at both mRNA and protein level was significantly upregulated in PHF2 KO myocytes compared to the CTRL (**Figure 4 D, E**). To validate whether PHF2 directly regulates *Gdi2* gene expression, we analyzed a previously published PHF2-ChIP sequencing dataset^26^ and detected a significant enrichment of PHF2 protein on *Gdi2* promoter (**Figure 4F**). We then performed a proximity ligation assay (PLA) to quantify the interaction between RAB8a and GDI2. As shown in **Figure 4G**, a significantly increase in the number of RAB8a/GDI2 complexes was observed in PHF2 KO myocytes, which is completely rescued by overexpressing PHF2^S655E^ mutant. Considering that RAB8a protein level is comparable between CTRL and PHF2 KO myocytes (**Figure S4G**), this result suggests that RAB8a is sequestered and maintained in its inactive form (GDP-bound) in the cytosol, indicating that the interaction between LDs and mitochondria is impaired. Transmission electron microscopy and confocal imaging confirmed that ablation of PHF2 caused a significantly decreased number of LD-mitochondria contact sites, which was rescued by overexpressing PHF2^S655E^ (**Figure 4H-I**). Altogether, these data demonstrate that AMPKα2-PHF2 signaling pathway is pivotal to regulate the interaction between LDs-mitochondria by repressing *Gdi2* expression and thus favoring the activation state of RAB8a protein.

## Discussion

The lysine demethylase PHF2 has earned significant attention for its involvement in diverse physiological and pathological processes across various organs^26,34–36^. Over the past decade, PHF2 has emerged as a multifunctional protein with an especially critical role in maintaining energy homeostasis^25,26^. A recent *in vitro* study has shown that while the absence of PHF2 disrupts the expression of genes related to myogenesis and sarcomere processes, skeletal muscle differentiation remains unaffected^37^. Our results provide the first evidence that PHF2’s enzymatic activity is indispensable, in the physiological context of regenerative myogenesis, for regulating lipid droplet metabolism during muscle stem cell fate.

Lipid droplets have recently been recognized as essential regulators of muscle stem cell fate during regenerative myogenesis^13^. However, the upstream factors driving LD dynamics during MuSC activation, proliferation, or myocyte fusion remain largely unexplored. Our findings establish PHF2 as a critical upstream regulator of LD metabolism in this context. Ablation of PHF2 leads to LD accumulation, which impairs mitochondrial function, resulting in a significant reduction in ATP production and a subsequent failure of myocytes fusion into multinucleated myotubes. This underscores the importance of PHF2 in maintaining the metabolic conditions necessary for successful myogenesis.

Furthermore, we demonstrate that PHF2 acts as part of the AMPKα2 metabolic signaling pathway, where its enzymatic activity is essential for LD turnover in myocytes. Depletion of AMPKα2 phenocopies the lipid droplet accumulation observed in the PHF2 KO model, highlighting their functional interplay. However, the rescue of LD accumulation via overexpression of a phospho-mimetic PHF2 mutant (PHF2^S655E^) does not restore fusion in AMPKα2 KO myocytes (data not shown), supporting that AMPKα2 has additional, PHF2-independent roles in myocyte fusion. This aligns with prior findings that AMPKα2 is crucial for myocyte fusion through the regulation of actin dynamics via BAIAP2 phosphorylation^30^.

AMPK, as a central metabolic sensor, plays a pivotal role in cellular energy regulation. Previous studies have pointed out its function in promoting the close interaction between lipid droplets and mitochondria, a process mediated by the triggering of RAB proteins in their GTP-bound state^8,32,38,39^. This interaction is critical for efficient energy utilization and metabolic homeostasis. In our study, we identify a novel AMPK-dependent metabolic signaling pathway that integrates epigenetic and metabolic regulation. Specifically, this study shows that AMPKα2 activates the epigenetic modifier PHF2, which likely represses the expression of *Gdi2*. This repression is crucial for the formation of the RAB8a-mediated tethering complex that bridges lipid droplets and mitochondria. By facilitating this interaction, PHF2 and AMPKα2 collaboratively provide efficient lipid droplet turnover and mitochondrial function, which are essential for energy production during myocyte fusion.

This newly characterized pathway underlines the intricate crosstalk between metabolic sensing and epigenetic regulation, highlighting the AMPKα2-PHF2 axis as a key regulator of cellular metabolic plasticity during regenerative myogenesis.

## Supporting information

Supplemental informations

## Acknowledgments

We thank C. Angleraux and P. Contard from the PBES (UMS3444) for animal breeding. We thank E. Devevre, S. Dussurgey and T. Deborde from the AniRA-Cytometry platform (UMS3444) for their expertise in cell sorting, and D. Ressnikoff, B. Chapuis and E. Errazuriz-Cerda (CIQLE imaging center, UMS3453) for their help with image and EM acquisition. We thank Shigeaki Kato and Prof. Yuuki Imai for providing the PHF2^fl/fl^ mice We thank C. Degletagne for support with single-cell RNA-seq. Funding sources: ANR (ANR-11-BSV2-0017) and AFM MyoNeurAlp strategic plan.

## Author contributions

I.S. designed the study. D.C. and I.S. conceived, performed, analyzed experiments and wrote the manuscript. S.M. and M.P. performed and analyzed experiments. W.J. performed transcriptomic analysis. A.K. and R.M. provided AMPKα2^-/-^ mouse model. L.S. provided fundings. All authors read, edited and approved the manuscript.

## Supplemental information

Figures S1–S4

## STAR Methods

### KEY RESOURCES TABLE

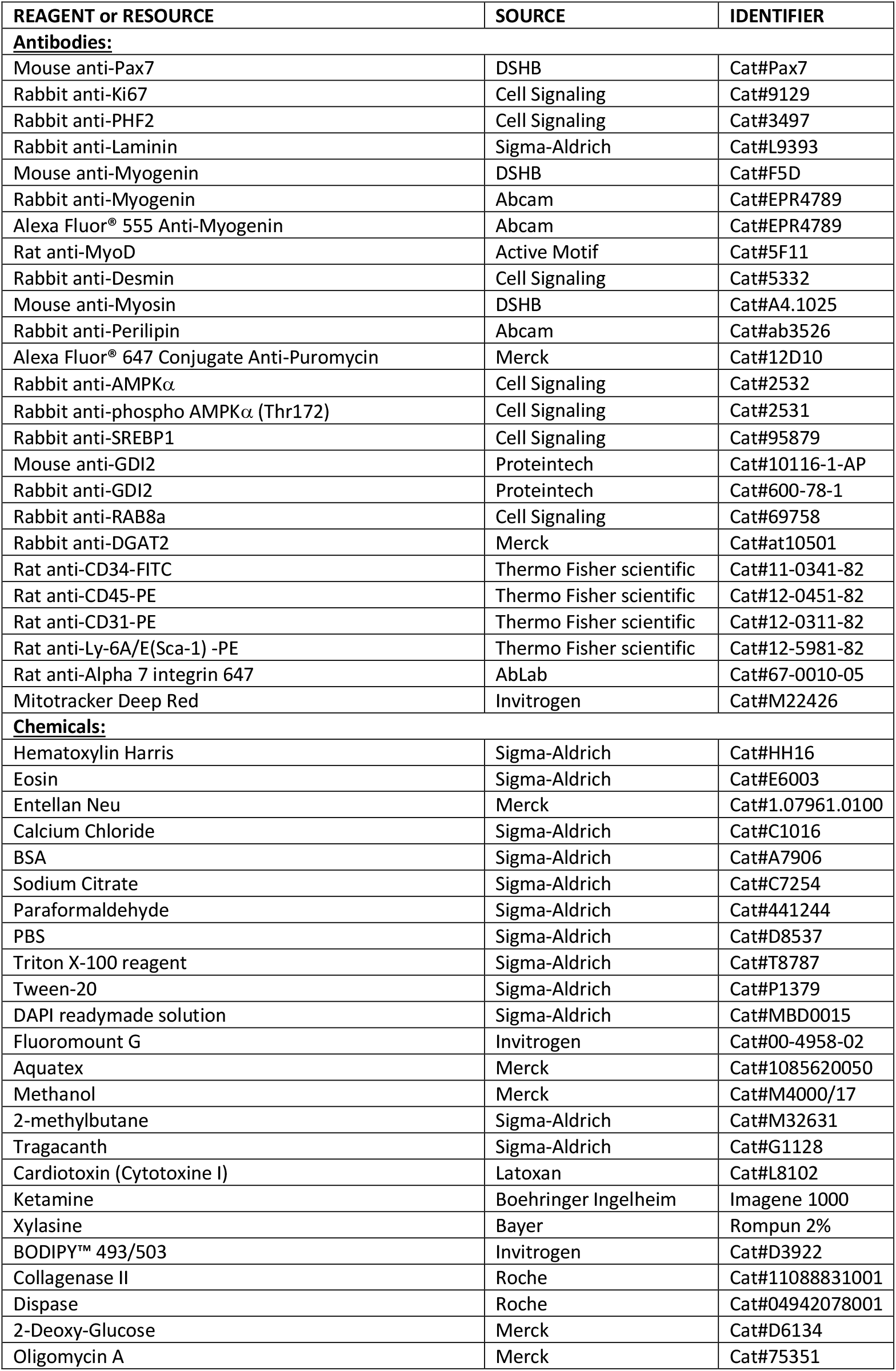
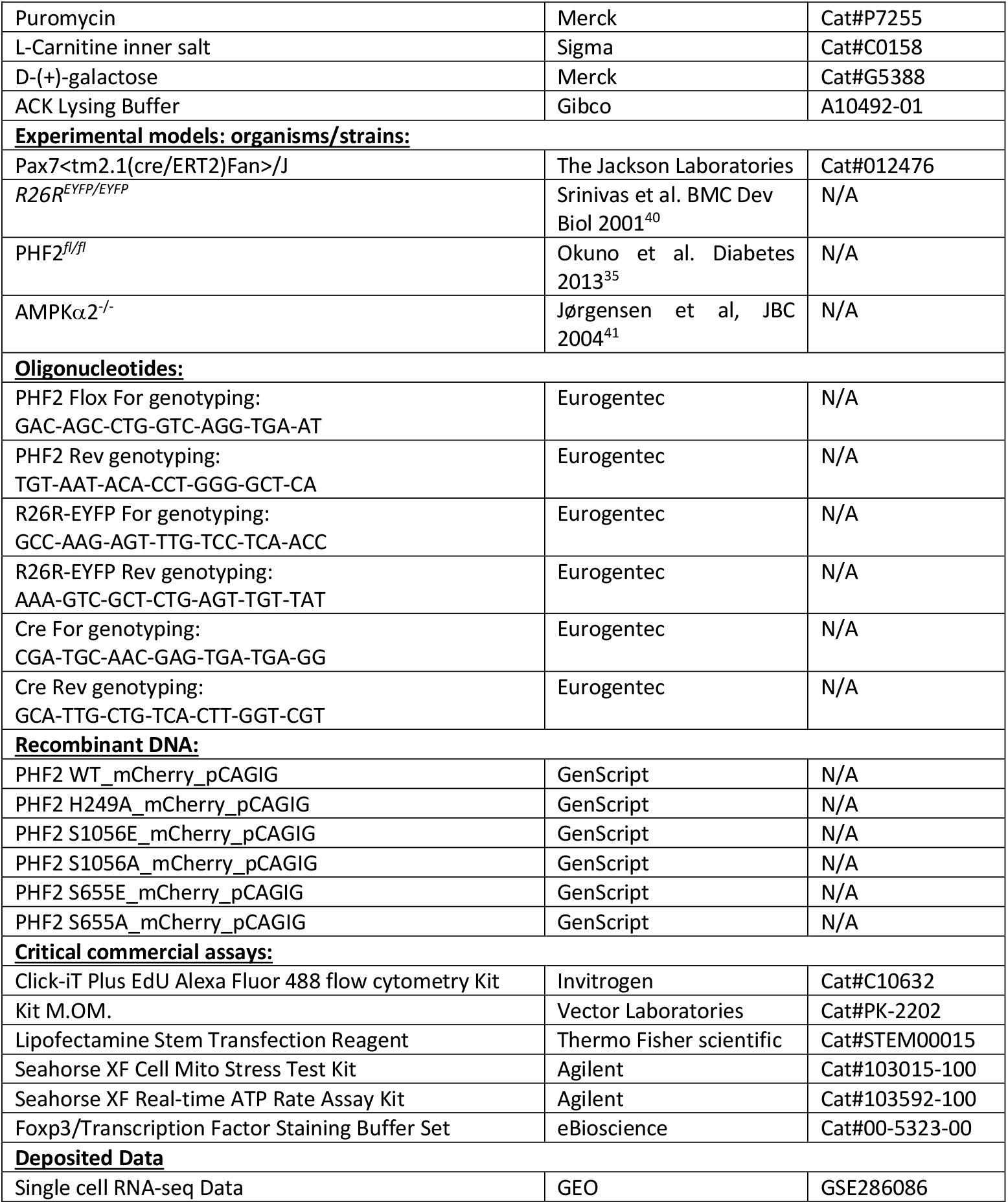

### Resource availability

Lead contact: Further information and requests for resources and reagents should be directed to and will be fulfilled by the lead contact, Isabella Scionti.

### Experimental model and subject details

#### Animals

All procedure on animals were performed in accordance with European regulations on animal experimentation and were approved by the local animal ethics committee (CECCAPP, University of Lyon) under the reference Apafis#16930 and Apafis#46209. Mice were bred and housed in AniRA-PBES animal facility. They were maintained in a temperature- and humidity-controlled facility with a 12h light/dark cycle, free access to water and standard rodent show.

Mice were genotyped with conventional PCR using standard conditions. Allelic recombination under the Pax7^CreERT2^ allele was induced by intraperitoneal injections of 2mg of Tamoxifen (TAM) for 5 days. Skeletal muscle injury was induced by an injection of 50 μl of cardiotoxin (10μM) into hindlimb TA muscle using 30G syringes under anesthesia induced by intraperitoneal injection with Ketamine (100mg/kg) and Xylazine (10mg/kg) in sterile saline solution.

#### Cell lines

Adult primary MuSCs were maintained on Matrigel-coated dishes in GM: Dulbecco’s modified Eagle’s medium F12 supplemented with 20% horse serum, 5 ng/mL fibroblast growth factor (FGF), and antibiotics. Primary MuSCs were differentiated into myotubes by replacing GM with a low serum medium containing 2% horse serum with antibiotics (LSM). For experiments in galactose medium, primary MuSCs were differentiated into myotubes by replacing GM with a glucose free LSM containing 10% horse serum, 10mM of galactose and antibiotics.

## Methods details

### MuSC isolation

Adult primary MuSCs were isolated from skeletal muscle tissue as previously described ^9^. Digested tissue was stained using antibodies: 0.2μg PE-conjugated rat anti-mouse CD31, 0.2μg PE-conjugated rat anti-mouse CD45, 0.2μg PE-conjugated rat anti-mouse Sca-1, 5μg 647-conjugated rat anti-mouse integrin alpha7 and 2.5μg rat anti-mouse CD34. Cells were incubated with primary antibodies for 40 minutes on ice. Adult primary MuSCs were FACS isolated into MuSC medium based on cell surface antigen markers: CD31-/CD45-/Sca1-/integrin-a7+/CD34+ using a FACsAria III.

### Cell treatments

For proliferation assay EdU solution (20mM) has added to adult primary MuSCs for 2 hours before the collection of cells. Cells were analyzed using flow cytometry (FACS Cantoll). For L-carnitine treatment, mononucleated cells were treated for 1 hour with 1mM L-carnitine inner salt dissolved in cell medium before the seahorse assay.

### Cell Transfection

Primary MuSCs were cultured in 24-well plates at 10.000 cells/cm^2^ in GM and, once adhered, transfected with 500ng of plasmids. After 24h, LSM was added for 48h, before performing immunofluorescence experiment.

### Immunoblotting

Protein extraction was performed from mononucleated cells 24h post LSM and quantified using the DC protein assay (Bio-Rad). Protein extracts were separated by electrophoresis on 8% and 10% SDS-PAGE and transferred onto polyvinylidene fluoride (PVDF) Immobilon-P membranes. Immunoblots were revealed with enhanced chemiluminescence (ECL) PLUS reagent according to the manufacturer’s instructions.

### Immunofluorescence

Permanox chamber slides (Nunc Lab-Tek) were used and cells were fixed with 4% (v/v) paraformaldehyde (PFA) for 10 minutes. Following fixation, material was permeabilized with 0.5% (v/v) Triton X-100 solution for 5 minutes and then blocked with 1% BSA, 0.2% Triton X-100 and 5% (v/v) goat serum for 60 minutes to reduce nonspecific antibody binding. Cells were then incubated with the following antibodies overnight at 4±C, 1:50 mouse anti-PAX7, 1:500 rabbit anti-Myogenin, 1:50 mouse anti-MyHC, 1:250 rat anti-MyoD, 1:200 rabbit anti-Desmin. Species-specific fluorochrome-conjugated secondary antibodies were then applied for 1 h at room temperature, before being mounted with 100 ng/ml of DAPI. Lipid droplets were stained with 1:1000 Bodipy 493/503 dye for 30 minutes at 37°C.

### Muscle Histology and immunohistochemistry and immunofluorescence

Mice were sacrificed and TA muscles were dissected and attached in Tragacanth gum, frozen in a cold 2-methylbutane bath and cryo-sectioned onto glass slides. Tissue sections were freshly fixed with PFA 4% for 10 minutes, washed three washes in PBS, permeabilized in methanol for 6 minutes at -20°C and then washed three times in PBS. After 10 minutes incubation in a hot antigen retrieval buffer, sections were washed three times in PBS, 0,1% triton X-100 (PBS-T) and then were saturated 2 hours at room temperature with M.O.M Mouse IGG Blocking reagent. The antigen retrieval buffer contained 10mM sodium citrate acid and 0,05% tween-20 and was adjusted at pH 6,0. Tissue sections were washed once in PBS-T, incubated for 5 minutes in M.O.M diluent prepared according to the manufacturer and stained at 4°C overnight with primary antibodies diluted in M.O.M diluent (1/200 for anti-KI67, 1/50 for anti-PHF2, 1/200 for anti-laminin, 1/50 for anti-Myogenin, 1/200 for anti-Perilipin, 1/50 for anti-PAX7). After three washes in PBS-T, sections were incubated for 1 hour at room temperature with secondary antibody and DAPI diluted in M.O.M diluent. After three washes, sections were mounted with Fluoromount-G. Fluorescent images were acquired on a Zeiss Z1 Axioscan. Hematoxylin and Eosin (H&E) staining was performed following standard methods.

### Seahorse mitochondrial respiration analysis

Mitochondrial respiration was measured with Seahorse XFe96 Analyzer (Agilent Technologies) according to the manual of Seahorse XF Cell Mito Stress Test Kit (Agilent Technologies). Briefly, MuSCs isolated by FACS were seeded for 6 days in GM and 2 days in LSM. After differentiation, myotubes were discarded after filtration (40 μm) and mononucleated cells were recovered and plated on Matrigel-coated XF96 cell culture microplate at a density of 8.000 cells per well overnight. Seahorse sensor cartridge was hydrated with calibrant in a non-CO2 incubator at 37°C for overnight 1 day before measurement. On the day of measurements, cells were washed twice and switched to Seahorse XF base medium (pH7.4, Agilent Technologies) supplemented with 1 mM sodium pyruvate, 1 mM L-glutamine, and 5 mM glucose (0mM glucose for Galactose analysis). Cells were equilibrated at 37°C in a non-CO2 incubator for 1 hour. The oxygen consumption rate for Mito Stress Test was monitored at the basal state and after sequential injection of the mitochondrial compounds oligomycin (1.5 μM), FCCP (2 μM), and Rotenone/antimycin A (1 μM) to induce mitochondrial stress. The ATP production rate for ATP Rate assay was monitored at the basal state and after sequential injection of the mitochondrial compounds oligomycin (1.5 μM), and Rotenone/antimycin A (1 μM) to induce mitochondrial stress. All mitochondrial respiration rates were generated and automatically calculated by the Seahorse Wave software with normalization to the number of cells/well.

### SCENITH Analysis

MuSCs isolated by FACS were seeded for 6 days in GM and 2 days in LSM. After differentiation, myotubes were discarded and mononucleated cells were recovered to perform SCENITH as described before 29.

### Proximity Ligation Assay

Duolink® in situ PLA reagents were used to detect the interaction between GDI2 and RAB8a and the manufacturer’s protocol was followed. Briefly, mononucleated cells 24h post differentiation were cultured on 8 well Permanox chamber slides (Nunc Lab-Tek). Cells were fixed, permeabilized and incubated overnight with mouse anti-GDI2 and rabbit anti-RAB8a. The following day, cells were incubated with secondary antibodies conjugated to oligonucleotides (PLA probe PLUS and PLA probe MINUS) for 1 h at 37°C. Afterward, ligation solution containing two oligonucleotides (that hybridize to the PLA probes) and Ligase was added for 30 min at 37°C. A closed circle is only formed if the two PLA probes are in close proximity. Finally, the closed circle was amplified using rolling-circle amplification reaction and the product was hybridized to fluorescently-labeled oligonucleotides. The fluorescent spots generated from positive interactions were quantified using a confocal microscope (Leica TCS SP5).

### Electron Microscopy

For immunogold project, cell cultures were fixed with 2% glutaraldehyde (EMS) in 0.1 M sodium cacodylate (pH 7.4) buffer at room temperature. After washing three times in 0.2 M sodium cacodylate buffer, cell cultures were post-fixed with 1% aqueous osmium tetroxide (EMS) at room temperature for 1 hour and contrasted with tea Oolong in cacodylate 0.2MpH7.4 (EM grade) for 1 hour at room temperature. After washing three time in 0.2Msodium cacodylate and 1 time in water, they were dehydrated in a graded series of ethanol at room temperature and embedded in Epon. After polymerization, ultrathin sections (100 nm) were cut on a UC7 (Leica) ultramicrotome and collected on Nickel grids 150 mesh. Immunogold labelling was performed by flotation the grids on drops of reactive media. A first step of unmaking the epitope is carried out with metaperiodate 12.5%. Sections were successively washed four times in water. Then, nonspecific sites were coated with 1% BSA and 1% normal goat serum in 50 mM Tris-HCl, pH 7.4 for 2h at RT. Thereafter, incubation was carried out overnight at 4°C in wet chamber with primary antibody (Myog 1/100). For the S655E mutant incubation was carried out overnight at 4°C in wet chamber with primary antibodies (Myog and Phf2 1/100). Sections were successively washed three times in 50 mM Tris-HCl, pH 7.4 and pH 8.2 at RT. Nonspecific sites were coated with 1% BSA and 1% normal goat serum in 50 mM Tris-HCl, pH 8.2 for 1h30 at RT. They were transferred in a wet chamber for 1h30 at RT in 1% BSA,50 mM Tris-HCl, pH 8.2 for 1h30 at RT, labelled with gold conjugated Secondary antibody (Goat anti Rabbit 10nm for Myog and Goat anti Rabbit 5nm for Phf2) (Aurion). Sections were successively washed three times in 50 Mm Tris-HCl pH8.2 and pH 7.4 and three times infiltrated distilled water. The immunocomplex was fixed by a wash in glutaraldehyde 4% for 3min. Sections were stained with lead citrate for 5min and observed with a transmission electron microscope JEOL 1400JEM (Tokyo, Japan) operating at 100kV equipped with a camera Orius 600 gatan and Digital Micrograph. This microscope is located at CIQLE (Centre d’Imagerie Quantitative Lyon Est), platform UCBL.

### Single cell RNASeq Analysis

Data demultiplexed in fastq format were checked with fastq and fastq_screen to control quality sequencing. Sequencing reads were processed with *CellRanger* (v.7.0.1) to align and count data for each sample with no-conservation of rRNA during process and intron-mode. Genome and transcriptome used is the 10x reference (refdata-gex-mm10-2020-A). We used *SoupX* (v.1.6.2) to remove contamination in cells. We determined Doublet barcode with *scDblfinder* on cluster-based approach and *dbr=0*.*1*. After quality control, we retained 21.284 cells for downstream analysis and detected 18068 gene transcripts. For the downstream treatment for integrative analysis, we used sample as variable. We used merge dataset and used *JoinLayers* from *Seurat* (V.5.0.1) and splitted merge dataset in function of group value of interest. We performed merged object with *SCTransform* with glmGamPoi method, *ncells = number of cells0*.*5* on 3000 common features between samples, PCA on SCT assay, Harmony integration with 50 dimensions used, (s)KNN graph build on harmony reduction matrix with all dimensions. We used Leiden clustering with igraph method. For the visualization, we used FFT-accelerated Interpolation-based t-SNE (FitSNE) with these parameters: a modification of initialization (first and second Harmony dimensions are divided by standard deviation of first Harmony dimension multiplied by 0.0001). Perplexity list used is a list starting 30 to number of cells in integrated dataset divided by 100 and learning rate = number of cells in integrated dataset divided by 12. All harmony dimensions used and 1000 iterations to run. To determine clusters, we integrated 2 methodologies: 1) manual based on known genes with a strong association with a cell identity and 2) *FindTransferAnchors* and *TransfertData* R function from Seurat by using McKellar et al 42 as reference dataset. Biomarkers are identified after *PrepSCTFindMarkers* on SCT assay and performed with FindAllMarkers with MAST test. We ranked biomarkers bases on p-adj. Integrative analysis identified 19 Seurat clusters. To define each cluster, we analyzed the expression of canonical cell type markers in resolution=0.8 that highlighted five hypergroups defined as: 1) Myofibers, 2) Muscle stem cells, 3) Proliferating muscle stem cells, 4) Myocytes, 5) Others For the pathways analysis, we used MSigDB reference C2 for mouse. Before calculation, we selected only gene sets sharing 75 % of genes in gene sets are in dataset. We used *AddModuloScore_Ucell* and *SmoothkNN* from *UCell* R package (v.2.6.2) on our dataset. All statistical view were made using *VlnPlot2* from *SeuratExtend* (v.1.0.0) and the statistic method is the non-parametric Wilcoxon test and Holm test post hoc.

## References

1. Madsen, S., Ramosaj, M., and Knobloch, M. (2021). Lipid metabolism in focus: how the build-up and breakdown of lipids affects stem cells. Development 148, dev191924. 10.1242/dev.191924.

2. Ryall, J.G., Cliff, T., Dalton, S., and Sartorelli, V. (2015). Metabolic Reprogramming of Stem Cell Epigenetics. Cell Stem Cell 17, 651–662. 10.1016/j.stem.2015.11.012.

3. Tatapudy, S., Aloisio, F., Barber, D., and Nystul, T. (2017). Cell fate decisions: emerging roles for metabolic signals and cell morphology. EMBO reports 18, 2105–2118. 10.15252/embr.201744816.

4. Cicciarello, D., and Scionti, I. (2023). [The unexpected role of lipid droplets in the regulation of muscle stem cells fate]. Med Sci (Paris) 39 Hors série n° 1, 28–31. 10.1051/medsci/2023144.

5. Farese, R.V., and Walther, T.C. (2009). Lipid Droplets Finally Get a Little R-E-S-P-E-C-T. Cell 139, 855–860. 10.1016/j.cell.2009.11.005.

6. Smith, B.K., Marcinko, K., Desjardins, E.M., Lally, J.S., Ford, R.J., and Steinberg, G.R. (2016). Treatment of nonalcoholic fatty liver disease: role of AMPK. American Journal of Physiology-Endocrinology and Metabolism 311, E730–E740. 10.1152/ajpendo.00225.2016.

7. Yang, J., Zhang, X., Yi, L., Yang, L., Wang, W.E., Zeng, C., Mi, M., and Chen, X. (2019). Hepatic PKA inhibition accelerates the lipid accumulation in liver. Nutr Metab (Lond) 16, 69. 10.1186/s12986-019-0400-5.

8. Li, Y., Munoz-Mayorga, D., Nie, Y., Kang, N., Tao, Y., Lagerwall, J., Pernaci, C., Curtin, G., Coufal, N.G., Mertens, J., et al. (2024). Microglial lipid droplet accumulation in tauopathy brain is regulated by neuronal AMPK. Cell Metab, S1550-4131(24)00118-9. 10.1016/j.cmet.2024.03.014.

9. Mouradian, S., Cicciarello, D., Lacoste, N., Risson, V., Berretta, F., Le Grand, F., Rose, N., Simonet, T., Schaeffer, L., and Scionti, I. (2024). LSD1 controls a nuclear checkpoint in Wnt/β-Catenin signaling to regulate muscle stem cell self-renewal. Nucleic Acids Res, gkae060. 10.1093/nar/gkae060.

10. Ryall, J.G., Dell’Orso, S., Derfoul, A., Juan, A., Zare, H., Feng, X., Clermont, D., Koulnis, M., Gutierrez-Cruz, G., Fulco, M., et al. (2015). The NAD(+)-dependent SIRT1 deacetylase translates a metabolic switch into regulatory epigenetics in skeletal muscle stem cells. Cell Stem Cell 16, 171–183. https://doi.org/mounier.

11. Theret, M., Gsaier, L., Schaffer, B., Juban, G., Ben Larbi, S., Weiss-Gayet, M., Bultot, L., Collodet, C., Foretz, M., Desplanches, D., et al. (2017). AMPKα1-LDH pathway regulates muscle stem cell self-renewal by controlling metabolic homeostasis. EMBO J 36, 1946–1962. 10.15252/embj.201695273.

12. Pala, F., Di Girolamo, D., Mella, S., Yennek, S., Chatre, L., Ricchetti, M., and Tajbakhsh, S. (2018). Distinct metabolic states govern skeletal muscle stem cell fates during prenatal and postnatal myogenesis. Journal of Cell Science 131, jcs212977. 10.1242/jcs.212977.

13. Yue, F., Oprescu, S.N., Qiu, J., Gu, L., Zhang, L., Chen, J., Narayanan, N., Deng, M., and Kuang, S. (2022). Lipid droplet dynamics regulate adult muscle stem cell fate. Cell Reports 38, 110267. 10.1016/j.celrep.2021.110267.

14. Wang, L., Liu, J., Miao, Z., Pan, Q., and Cao, W. (2021). Lipid droplets and their interactions with other organelles in liver diseases. The International Journal of Biochemistry & Cell Biology 133, 105937. 10.1016/j.biocel.2021.105937.

15. Hugenroth, M., and Bohnert, M. (2020). Come a little bit closer! Lipid droplet-ER contact sites are getting crowded. Biochimica et Biophysica Acta (BBA) - Molecular Cell Research 1867, 118603. 10.1016/j.bbamcr.2019.118603.

16. Bosch, M., Parton, R.G., and Pol, A. (2020). Lipid droplets, bioenergetic fluxes, and metabolic flexibility. Seminars in Cell & Developmental Biology 108, 33–46. 10.1016/j.semcdb.2020.02.010.

17. Olzmann, J.A., and Carvalho, P. (2019). Dynamics and functions of lipid droplets. Nat Rev Mol Cell Biol 20, 137–155. 10.1038/s41580-018-0085-z.

18. Wanders, R.J.A., Waterham, H.R., and Ferdinandusse, S. (2016). Metabolic Interplay between Peroxisomes and Other Subcellular Organelles Including Mitochondria and the Endoplasmic Reticulum. Front Cell Dev Biol 3, 83. 10.3389/fcell.2015.00083.

19. Horton, J.R., Upadhyay, A.K., Hashimoto, H., Zhang, X., and Cheng, X. (2011). Structural basis for human PHF2 Jumonji domain interaction with metal ions. J Mol Biol 406, 1–8. 10.1016/j.jmb.2010.12.013.

20. Baba, A., Ohtake, F., Okuno, Y., Yokota, K., Okada, M., Imai, Y., Ni, M., Meyer, C.A., Igarashi, K., Kanno, J., et al. (2011). PKA-dependent regulation of the histone lysine demethylase complex PHF2–ARID5B. Nat Cell Biol 13, 668–675. 10.1038/ncb2228.

21. Dong, Y., Hu, H., Zhang, X., Zhang, Y., Sun, X., Wang, H., Kan, W., Tan, M., Shi, H., Zang, Y., et al. (2023). Phosphorylation of PHF2 by AMPK releases the repressive H3K9me2 and inhibits cancer metastasis. Sig Transduct Target Ther 8, 1–17. 10.1038/s41392-022-01302-6.

22. Okuno, Y., Ohtake, F., Igarashi, K., Kanno, J., Matsumoto, T., Takada, I., Kato, S., and Imai, Y. (2013). Epigenetic regulation of adipogenesis by PHF2 histone demethylase. Diabetes 62, 1426–1434. 10.2337/db12-0628.

23. Lepper, C., Conway, S.J., and Fan, C.-M. (2009). Adult satellite cells and embryonic muscle progenitors have distinct genetic requirements. Nature 460, 627–631. 10.1038/nature08209.

24. Saclier, M., Yacoub-Youssef, H., Mackey, A.L., Arnold, L., Ardjoune, H., Magnan, M., Sailhan, F., Chelly, J., Pavlath, G.K., Mounier, R., et al. (2013). Differentially activated macrophages orchestrate myogenic precursor cell fate during human skeletal muscle regeneration. Stem Cells 31, 384–396. 10.1002/stem.1288.

25. Jeong, D.-W., Park, J.-W., Kim, K.S., Kim, J., Huh, J., Seo, J., Kim, Y.L., Cho, J.-Y., Lee, K.-W., Fukuda, J., et al. (2023). Palmitoylation-driven PHF2 ubiquitination remodels lipid metabolism through the SREBP1c axis in hepatocellular carcinoma. Nat Commun 14, 6370. 10.1038/s41467-023-42170-0.

26. Bricambert, J., Alves-Guerra, M.-C., Esteves, P., Prip-Buus, C., Bertrand-Michel, J., Guillou, H., Chang, C.J., Vander Wal, M.N., Canonne-Hergaux, F., Mathurin, P., et al. (2018). The histone demethylase Phf2 acts as a molecular checkpoint to prevent NAFLD progression during obesity. Nat Commun 9, 2092. 10.1038/s41467-018-04361-y.

27. Relaix, F., Bencze, M., Borok, M.J., Der Vartanian, A., Gattazzo, F., Mademtzoglou, D., Perez-Diaz, S., Prola, A., Reyes-Fernandez, P.C., Rotini, A., et al. (2021). Perspectives on skeletal muscle stem cells. Nat Commun 12, 692. 10.1038/s41467-020-20760-6.

28. Marroquin, L.D., Hynes, J., Dykens, J.A., Jamieson, J.D., and Will, Y. (2007). Circumventing the Crabtree effect: replacing media glucose with galactose increases susceptibility of HepG2 cells to mitochondrial toxicants. Toxicol Sci 97, 539–547. 10.1093/toxsci/kfm052.

29. Argüello, R.J., Combes, A.J., Char, R., Gigan, J.-P., Baaziz, A.I., Bousiquot, E., Camosseto, V., Samad, B., Tsui, J., Yan, P., et al. (2020). SCENITH: A Flow Cytometry-Based Method to Functionally Profile Energy Metabolism with Single-Cell Resolution. Cell Metab 32, 1063-1075.e7. 10.1016/j.cmet.2020.11.007.

30. Kneppers, A., Ben Larbi, S., Theret, M., Saugues, A., Dabadie, C., Gsaier, L., Ferry, A., Rhein, P., Gondin, J., Sakamoto, K., et al. (2023). AMPKα2 is a skeletal muscle stem cell intrinsic regulator of myonuclear accretion. iScience 26, 108343. 10.1016/j.isci.2023.108343.

31. Benador, I.Y., Veliova, M., Liesa, M., and Shirihai, O.S. (2019). Mitochondria Bound to Lipid Droplets: Where Mitochondrial Dynamics Regulate Lipid Storage and Utilization. Cell Metabolism 29, 827–835. 10.1016/j.cmet.2019.02.011.

32. Ouyang, Q., Chen, Q., Ke, S., Ding, L., Yang, X., Rong, P., Feng, W., Cao, Y., Wang, Q., Li, M., et al. (2023). Rab8a as a mitochondrial receptor for lipid droplets in skeletal muscle. Developmental Cell 58, 289-305.e6. 10.1016/j.devcel.2023.01.007.

33. Müller, M.P., and Goody, R.S. (2017). Molecular control of Rab activity by GEFs, GAPs and GDI. Small GTPases 9, 5. 10.1080/21541248.2016.1276999.

34. Lee, K.-H., Ju, U.-I., Song, J.-Y., and Chun, Y.-S. (2014). The histone demethylase PHF2 promotes fat cell differentiation as an epigenetic activator of both C/EBPα and C/EBPδ. Mol Cells 37, 734–741. 10.14348/molcells.2014.0180.

35. Okuno, Y., Ohtake, F., Igarashi, K., Kanno, J., Matsumoto, T., Takada, I., Kato, S., and Imai, Y. (2013). Epigenetic regulation of adipogenesis by PHF2 histone demethylase. Diabetes 62, 1426–1434. 10.2337/db12-0628.

36. Kim, H.-J., Park, J.-W., Lee, K.-H., Yoon, H., Shin, D.H., Ju, U.-I., Seok, S.H., Lim, S.H., Lee, Z.H., Kim, H.-H., et al. (2014). Plant homeodomain finger protein 2 promotes bone formation by demethylating and activating Runx2 for osteoblast differentiation. Cell Res 24, 1231–1249. 10.1038/cr.2014.127.

37. Fukushima, T., Hasegawa, Y., Kuse, S., Fujioka, T., Nikawa, T., Masubuchi, S., and Sakakibara, I. (2024). PHF2 regulates sarcomeric gene transcription in myogenesis. PLOS ONE 19, e0301690. 10.1371/journal.pone.0301690.

38. Herms, A., Bosch, M., Reddy, B.J.N., Schieber, N.L., Fajardo, A., Rupérez, C., Fernández-Vidal, A., Ferguson, C., Rentero, C., Tebar, F., et al. (2015). AMPK activation promotes lipid droplet dispersion on detyrosinated microtubules to increase mitochondrial fatty acid oxidation. Nat Commun 6, 7176. 10.1038/ncomms8176.

39. Samovski, D., Su, X., Xu, Y., Abumrad, N.A., and Stahl, P.D. (2012). Insulin and AMPK regulate FA translocase/CD36 plasma membrane recruitment in cardiomyocytes via Rab GAP AS160 and Rab8a Rab GTPase. Journal of Lipid Research 53, 709–717. 10.1194/jlr.M023424.

40. Srinivas, S., Watanabe, T., Lin, C.-S., William, C.M., Tanabe, Y., Jessell, T.M., and Costantini, F. (2001). Cre reporter strains produced by targeted insertion of EYFP and ECFP into the ROSA26 locus. BMC Dev Biol 1, 4. 10.1186/1471-213X-1-4.

41. Jørgensen, S.B., Viollet, B., Andreelli, F., Frøsig, C., Birk, J.B., Schjerling, P., Vaulont, S., Richter, E.A., and Wojtaszewski, J.F.P. (2004). Knockout of the α2 but Not α1 5’-AMP-activated Protein Kinase Isoform Abolishes 5-Aminoimidazole-4-carboxamide-1-β-4-ribofuranosidebut Not Contraction-induced Glucose Uptake in Skeletal Muscle*. Journal of Biological Chemistry 279, 1070–1079. 10.1074/jbc.M306205200.

42. McKellar, D.W., Walter, L.D., Song, L.T., Mantri, M., Wang, M.F.Z., De Vlaminck, I., and Cosgrove, B.D. (2021). Large-scale integration of single-cell transcriptomic data captures transitional progenitor states in mouse skeletal muscle regeneration. Commun Biol 4, 1280. 10.1038/s42003-021-02810-x.

