## Supplemental informations for "The AMPKα2/PHF2 axis is critical for turning over lipid droplets during muscle stem cell fate"

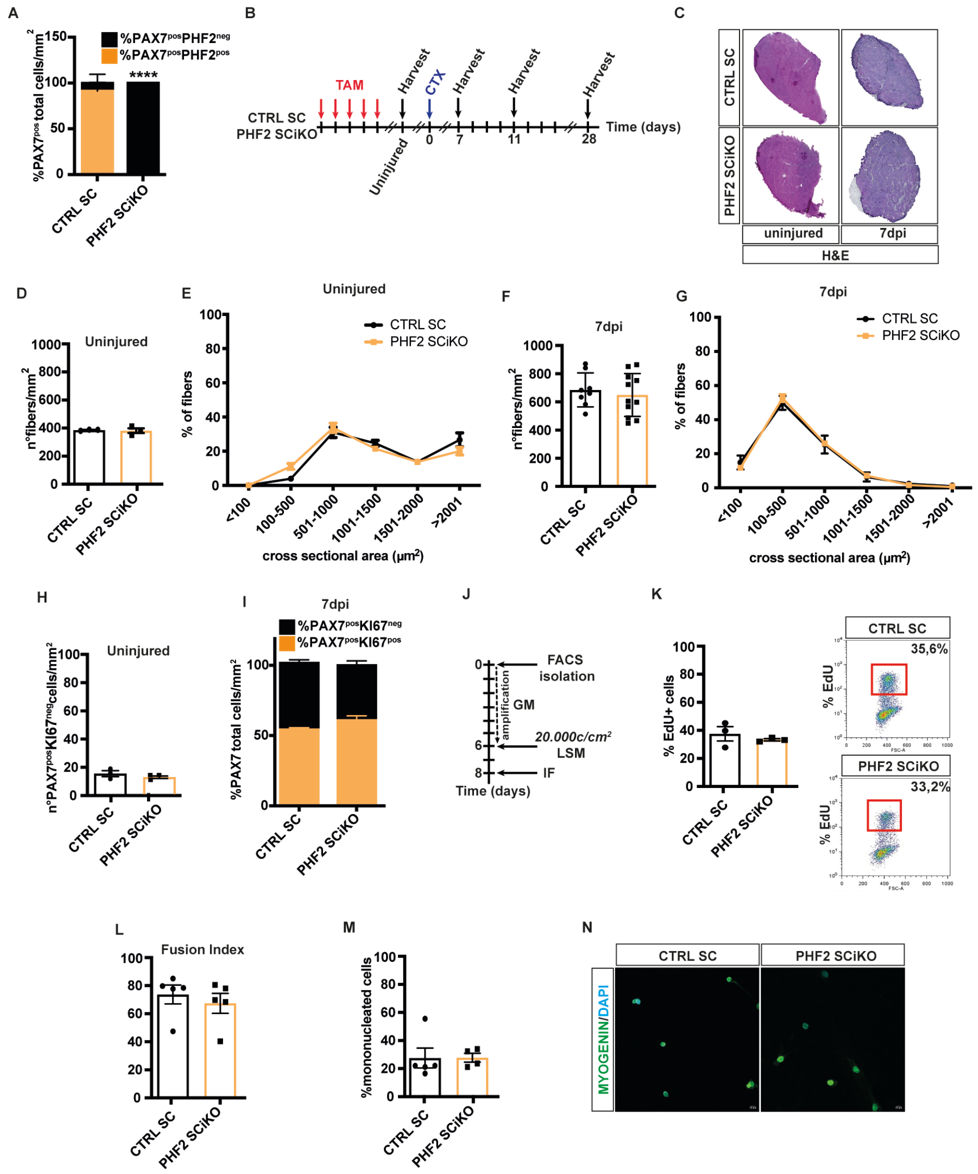

**Figure S1: PHF2 is essential to ensure muscle stem cell function and skeletal muscle regeneration *in vivo*, related to Figure 1.**

(A) Quantification of the number of PAX7$^{pos}$PHF2$^{neg}$ cells per mm$^2$ in uninjured CTRL SC and PHF2 SCiKO TA cryosections. (B) CTX experimental setup. (C) Hematoxylin-eosin staining on cryosections of TA in uninjured and 7dpi. Quantification of the number of myofibers per mm$^2$ (D) and CSA distribution of muscle fibers (E) in uninjured TA muscles. Quantification of the number of myofibers per mm$^2$ (F) and CSA distribution of muscle fibers (G) of regenerating TA muscles at 7dpi. (H) Quantification of the number of sublaminar PAX7$^{pos}$KI67$^{neg}$ cells per mm$^2$ in uninjured TA cryosections. (I) Percentage of PAX7$^{pos}$KI67$^{pos}$ MuSCs (orange) per mm$^2$ at 7dpi. (J) Experimental setup. (K) Percentage of CTRL SC and PHF2 SCiKO MuSCs in S-Phase. Fusion Index (L) and percentage of mononucleated cells (M) of FACS-isolated MuSCs after two days in LSM. (N) Representative images of committed myocytes (MYOG$^{pos}$). n = 3-11 mice/genotype. n = 3-5 primary MuSC cultures/genotype. Values are mean or percentage mean ± SEM. **** P < 0.0001 (Tukey's test after one way-ANOVA [A panel]).

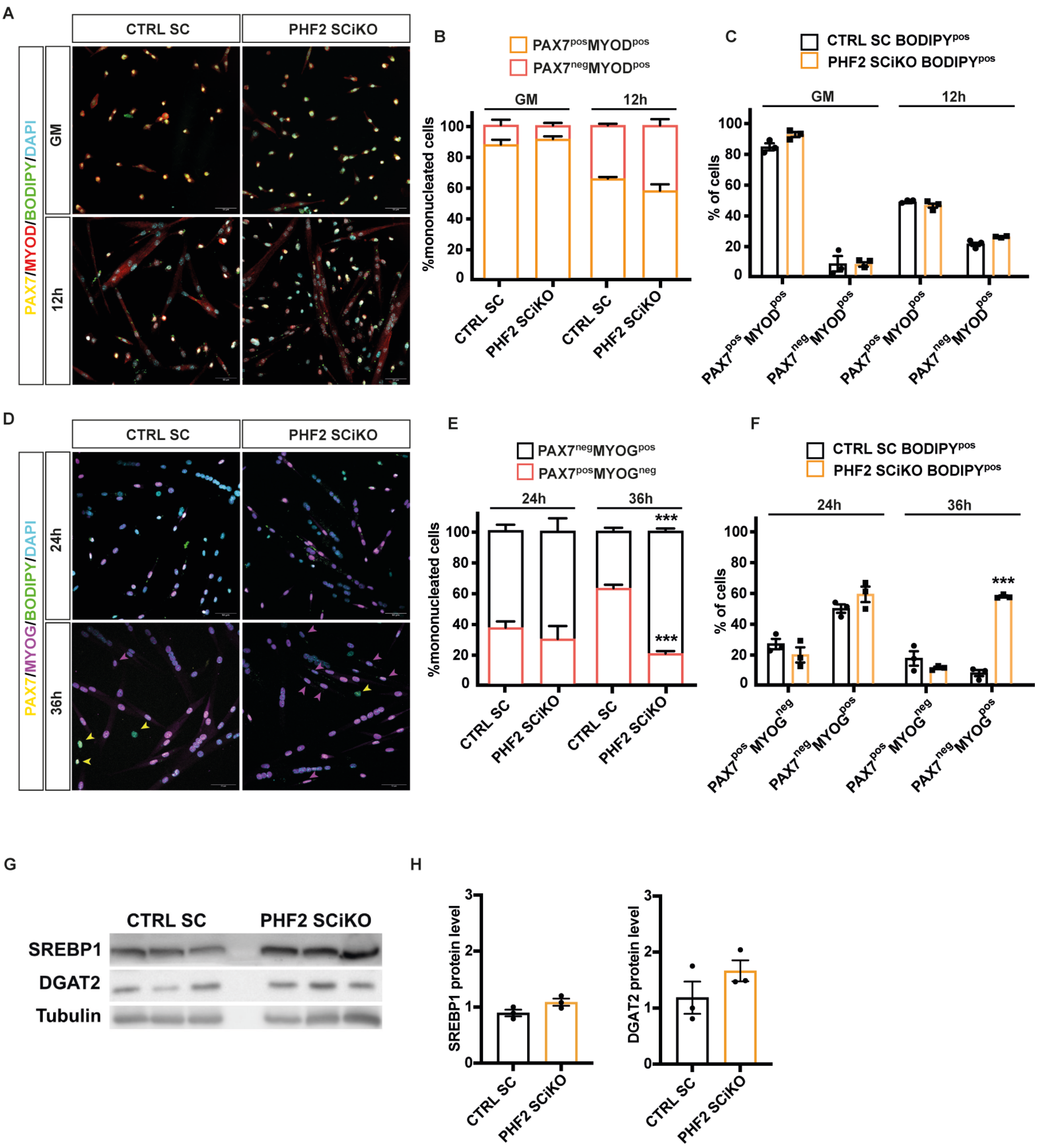


**Figure S2: PHF2 depleted committed myocytes starts to accumulate lipid droplets 36h post differentiation, related to Figure 2.**

Representative images (A) of the percentage of proliferating myoblast (PAX7$^{pos}$MYOD$^{pos}$) and committed myoblasts (PAX7$^{neg}$MYOD$^{pos}$) (B) which accumulate lipid droplets (BODIPY$^{pos}$) (C) in proliferation (GM) and 12h in low serum medium. Representative images (D) of the percentage of differentiated myocytes (purple arrowhead, PAX7$^{neg}$MYOG$^{pos}$) and PAX7$^{pos}$MYOG$^{neg}$ (yellow arrowhead) (E), which accumulate lipid droplets (BODIPY$^{pos}$) (F) at 24h and 36h in LSM. (G) Immunoblot of SREBP1 and DGAT2 proteins and (H) quantification of the immunoblot, tubulin was used as loading control. Scale bars, 50 µm. n = 3 primary MuSC cultures/genotype. Values are mean or percentage mean ± SEM. ***P < 0.001 (Student *t*-test [B-C-E-F-H panels]).

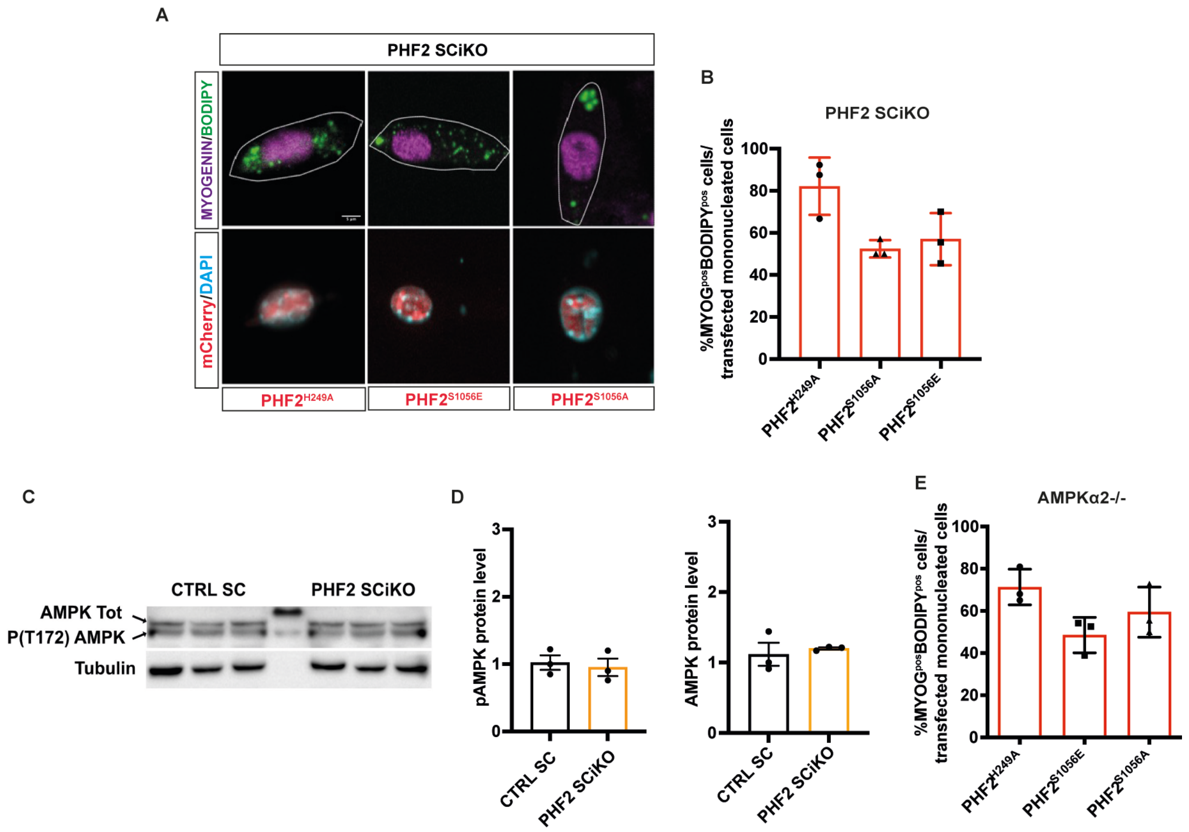


**Figure S3: PKA-mediated phosphorylation of PHF2 is not required to regulate lipid droplet turnover in committed myocytes, related to Figure 3.**

Representative images (A) of the percentage of PHF2 SCiKO MYOG$^{pos}$BODIPY$^{pos}$ mononucleated cells transfected with either MOCK, PHF2$^{H249A}$ or PHF2$^{S1056E}$ and PHF2$^{S1056A}$ (B). (C) Immunoblot of AMPK and pAMPK proteins and (D) quantification of the immunoblot, tubulin was used as loading control. (E) Percentage of AMPK$\alpha$2-/- MYOG$^{pos}$BODIPY$^{pos}$ mononucleated cells transfected with either PHF2$^{H249A}$, PHF2$^{S1056E}$ or PHF2$^{S1056A}$. Scale bars, 5 µm. n = 3 primary MuSC cultures/genotype. Values are mean or percentage mean ± SEM.

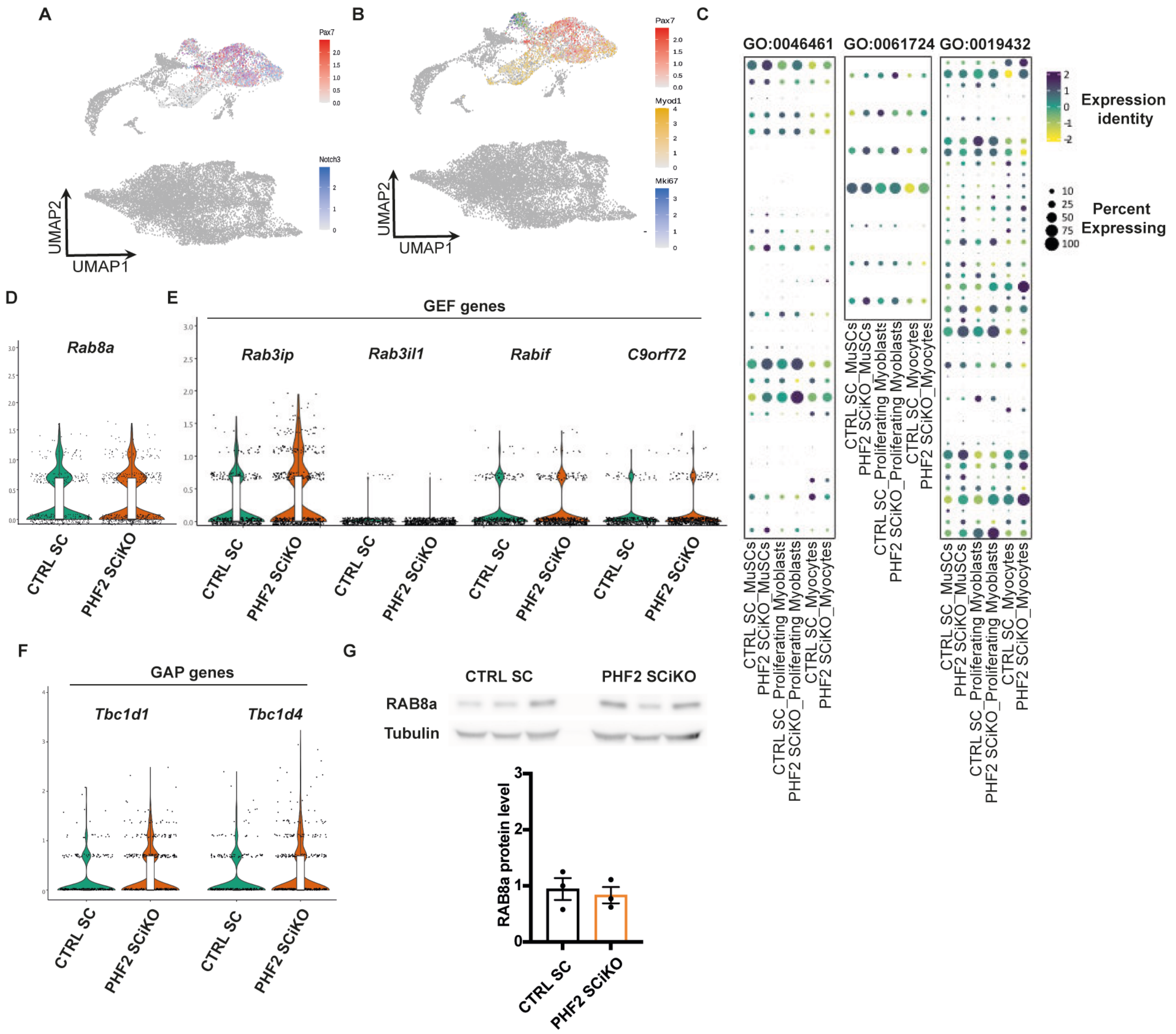


**Figure S4: PHF2 depletion does not perturb lipid metabolism gene's expression, related to Figure 4.**

MuSCs (A) and proliferating myoblasts (B) clusters. Expressions of lipolysis, lypophagy and LD biogenesis genes in CTRL SC and PHF2 SCiKO clusters (C). Violin plots of *Rab8a* (D), GEF genes (*Rab3ip, Rab3il1, Rabif and C9orf72*) (E) and GAP genes (*Tbc1d1* and *Tbc1d4*) (F) in CTRL SC and PHF2 SCiKO myocyte cluster. (G) RAB8a protein level and quantification in CTRL SC and PHF2 SCiKO mononucleated cells 24h post LSM. n = 3 primary MuSC cultures/genotype. Values are mean or percentage mean ± SEM.
